# Probing nanomechanics by direct indentation using Nanoendoscopy-AFM reveals the nuclear elasticity transition in cancer cells

**DOI:** 10.1101/2024.12.13.626302

**Authors:** Takehiko Ichikawa, Yohei Kono, Makiko Kudo, Takeshi Shimi, Naoyuki Miyashita, Tomohiro Maesaka, Kojiro Ishibashi, Kundan Sivashanmugan, Takeshi Yoshida, Keisuke Miyazawa, Rikinari Hanayama, Eishu Hirata, Kazuki Miyata, Hiroshi Kimura, Takeshi Fukuma

## Abstract

The assessment of nuclear structural changes is considered a potential biomarker of metastatic cancer. However, accurately measuring nuclear elasticity remains challenging. Traditionally, nuclear elasticity has been measured by indenting the cell membrane with a bead-attached atomic force microscopy (AFM) probe or aspirating isolated nuclei with a micropipette tip. However, indentation using a bead-attached probe is influenced by the cell membrane and cytoskeleton, while measurements of isolated nuclei do not reflect their intact state. In this study, we used Nanoendoscopy-AFM, a technique in which a nanoneedle probe is inserted into a living cell to directly measure nuclear elasticity and map its distribution. Our findings show that nuclear elasticity increases under serum depletion but decreases when serum-depleted cells are treated with TGF-β, which induces epithelial-mesenchymal transition (EMT). Furthermore, we found that changes in nuclear elasticity correlate positively with trimethylation levels of histone H4 at lysine 20, rather than with nuclear lamins expression levels. These findings suggest that alterations in chromatin structure underlie changes in nuclear elasticity during cancer progression.

## Introduction

Most solid tumors are detected through palpation, X-ray radiography, and ultrasound imaging due to their increased stiffness compared to surrounding tissue^1–3^. This stiffness is attributed to the stiffening of the surrounding stroma, which consists of the extracellular matrix (ECM) and supporting connective tissue cells. Stromal stiffening is believed to contribute to tumor rigidity and progression, as well as promote various aspects of cancer development^4–6^. Conversely, numerous studies on the elasticity of individual cancer cells suggest that cancer progression leads to cellular softening^7–9^. This softening is accompanied by the remodeling of cell-cell and cell-ECM interactions and is thought to enhance cell deformability, facilitating cancer cell migration through tight spaces^10^.

The transition in cell elasticity is influenced by the mechanical properties of the nucleus, cytoskeleton, and plasma membrane^11–13^. Specifically, nuclear elasticity has been investigated as a potential indicator of malignant cancer^14,15^. Measurements of nuclear elasticity in cervical, breast, and thyroid cancer cells have shown that cancer cell nuclei are softer than those of normal cells^16–18^. However, traditional methods may not accurately assess intact nuclear elasticity, as measurements are typically conducted by either indenting the cell membrane above a nucleus using a colloidal probe cantilever in atomic force microscopy (AFM) or measuring isolated nuclei using AFM or microaspiration^19–33^. The former method does not exclusively assess nuclear elasticity but also includes contributions from the cell membrane and cytoskeleton. Meanwhile, the latter method does not necessarily reflect the properties of an intact nucleus. Additionally, methods utilizing magnetic beads or optical tweezers cannot apply large forces and may fail to measure elasticity accurately^34,35^. Therefore, accurately measuring intact nuclear elasticity remains challenging, highlighting the need for a new and more precise method.

In mammalian cells, DNA wraps around histone octamers, forming nucleosomes, which are further organized into a higher-order chromatin structure and enclosed by the nuclear envelope. Chromatin can be divided into heterochromatin (tightly packed) and euchromatin (loosely packed)^36^. Heterochromatin is characterized by specific modifications, such as histone H3 lysine 9 dimethylation and trimethylation (H3K9me2 and H3K9me3, respectively), lysine 27 trimethylation (H3K27me3), and H4 lysine 20 trimethylation (H4K20me3), which mark transcriptionally inactive regions^37–40^. A decreased level of H3K9me3 and depletion of heterochromatin protein 1 (HP1) α cause nuclear softening^41^.

The nuclear envelope consists of double phospholipid bilayers—the inner and outer nuclear membranes (INM and ONM)—and associated proteins^42^. A part of heterochromatin is anchored to the nuclear lamina (NL) lining the INM. The NL is connected to the cytoskeletal system through the linker of the nucleoskeleton and cytoskeleton (LINC) complex, which is localized to the IMN and OMN and the lumen between them^43^. The major structural components of the NL are nuclear lamins, classified into A-type lamins (lamins A and C) and B-type lamins (lamins B1 and B2)^44,45^. These lamins assemble into a filament to form meshworks of the NL^46–49^. The NL is considered the primary determinant of nuclear mechanics^22,50^.

Recent advances in AFM technology have demonstrated that AFM with a nanoneedle probe can measure the mechanical properties of the nucleus in living cells. Nanoneedle-based AFM technology was first developed by Nakamura and colleagues and has been applied to molecular delivery and detection inside living cells^51,52^. Subsequently, Sun’s group showed that intact nuclear elasticity could be measured inside a living cell using a nanoneedle probe^53,54^. In that study, nuclear elasticity was estimated from force curves obtained once or several times without precise control over the position of the cell nucleus. A major limitation of this approach is the difficulty in detecting and eliminating the influence of obstacles, such as the cytoskeleton, vesicles, and assembled structures between the nuclear membrane and the cell membrane, which compromises the accuracy of nuclear elasticity measurements. This is a critical issue because the primary advantage of the nanoneedle-based AFM technology is its ability to measure intact nuclear elasticity while minimizing the influence of non-nuclear components, thereby providing an accurate assessment of nuclear elasticity.

To address this issue, we developed a method named “Nanoendoscopy-AFM (NE-AFM), ” which allows thousands of probe insertions into the cell without causing severe damage^55–57^. By optimizing the tip shape, refining insertion conditions, and developing dedicated analysis software, NE-AFM enables nanoscale mapping of nuclear surface elasticity. This technique improves the accuracy of nuclear elasticity measurement by selecting force curves that specifically reflect interactions with the nuclear membrane from a large dataset. In this study, we employed NE-AFM to precisely measure nuclear elasticity in living cancer cells with different metastatic potentials, exploring the relationship between nuclear mechanics and cancer metastasis.

## Results

### Whole-cell measurement using NE-AFM

Fig. 1a illustrates the process of whole-cell measurement using 3D NE-AFM. This technique repeatedly inserts a nanoneedle probe into the cell to measure the force versus distance (*F-z*) at arrayed-*xy* positions across the target area. The nanoneedle probe was fabricated by electron beam deposition (EBD) on the truncated tip of a commercial cantilever (BL-AC40TS-C2, spring constant: 0.1 N/m), as shown in Fig. 1b. The nanoneedle length of 5 μm is long enough to measure the elasticity of the nuclear surface inside the living cell. The diameter must be less than 200 nm because a thicker nanoneedle probe damages the cell^55^. In this experiment, the diameter was controlled to be around 160 nm, which is well below this threshold value (Fig. 1b).

**Fig. 1.**
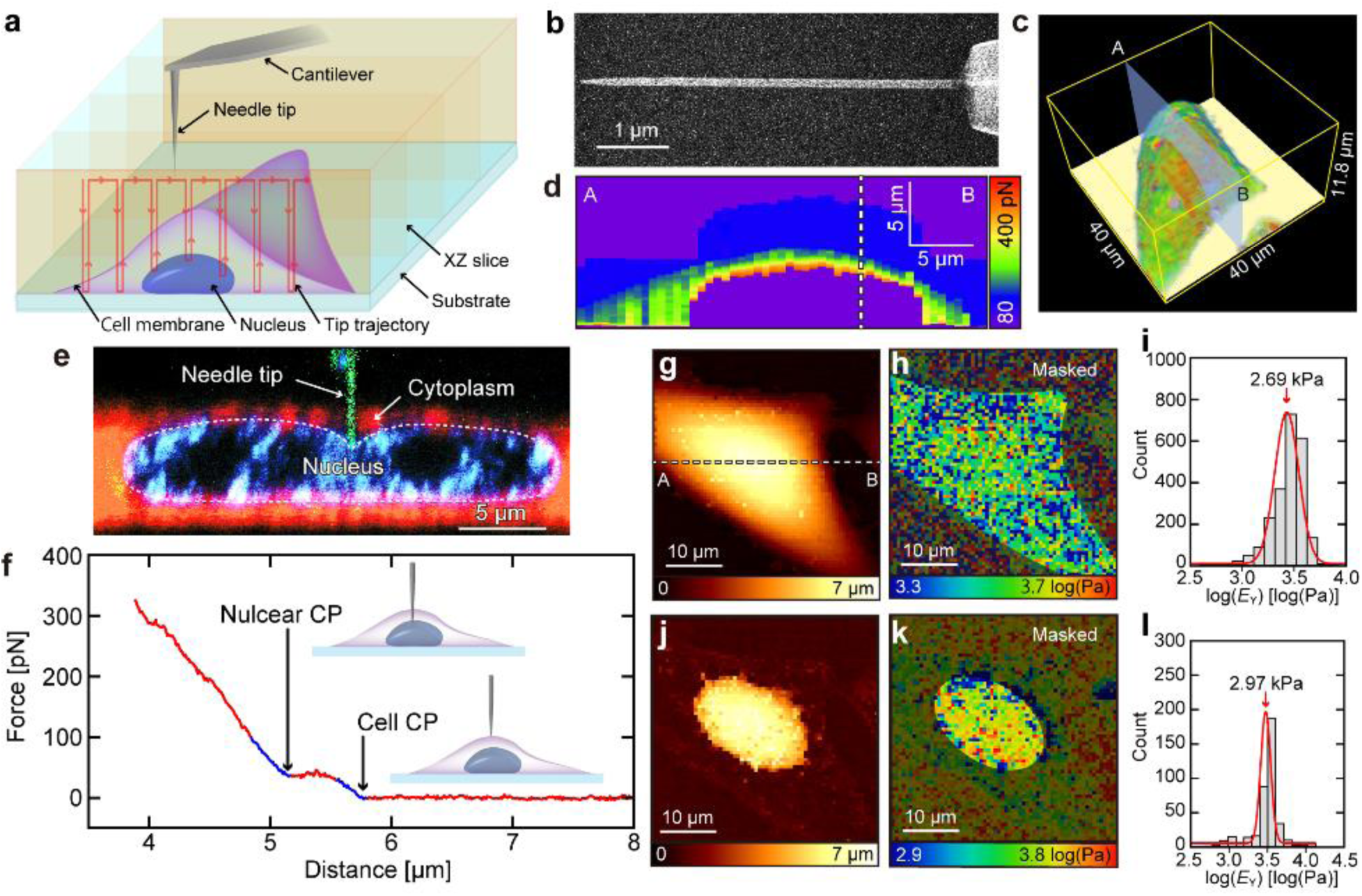
Nanoendoscopy-AFM (NE-AFM) measurement of a whole cell. (a) Schematic illustration of the NE-AFM applied to a whole cell. (b) Scanning electron microscope image of nanoneedle probe. (c) Volume rendering of the obtained 3D force map with a force-weighted transparency filter to visualize cell membrane surfaces. (d) A cross-sectional force map was taken along line AB in panel (c) and (g). (e) Cross-sectional image showing the tip pressing against the nucleus. The nanoneedle probe, cytoplasm, and nucleus were stained green, red, and blue, respectively. (f) Typical force-distance (*F-z*) curve during tip approach. The force represents the force applied to the tip, while the distance indicates the relative tip height with respect to the arbitrarily determined zero position. Cell contact point (Cell CP) and nuclear contact point (Nuclear CP) are marked. (g) Cell CP map. (h) Young’s modulus (*E*_Y_) map of the cell membrane. (i) Histogram of *E*_Y_ values for the cell membrane (mean value of the Gaussian fit: 2.69 kPa). (j) Nuclear CP map. (k) *E*_Y_ map of the nuclear surface. (l) Histogram of *E*_Y_ values for the nuclear surface (mean value of the Gaussian fit: 2.97 kPa).

Fig. 1c presents a volume-rendered cell surface image constructed from the obtained 3D force map with a force-weighted transparency filter. Fig. 1d depicts the cross-sectional force map along the line A-B in Figs. 1c and g. A typical *F-z* curve is displayed in Fig. 1f, where “distance ” refers to the relative tip height with respect to the arbitrarily determined zero position, and “force ” represents the vertical force applied to the tip. The local rise around 5.7 µm in the force curve (right blue line part) indicates the initial contact point with the cell membrane (cell CP), showing a plateau after membrane penetration. Another rise, at approximately 5 µm (left blue line part), marks the contact point with the nucleus (nuclear CP). Fig. 1e displays the cross-sectional fluorescence image of the deformed nucleus during its indentation by needle probe. By fitting the Herz-Sneddon model to each indentation profile in the force curve, Young’s moduli (*E*_Y_) of the cell membrane and the nucleus can be estimated separately^58^.

By extracting data from the 3D force map, we can reconstruct the cell membrane’s CP and elasticity maps (Figs. 1g and h). Fig. 1i presents a histogram of the cell membrane’s *E*_Y_ (peak value: 2,693 Pa). Similarly, we can reconstruct nuclear CP and elasticity maps (Figs. 1j and k), with Fig. 1l showing the *E*_Y_ histogram for the nucleus (peak value: 2,972 Pa). Analyses were performed using custom software developed in-house^57^.

### Measurement of the nuclear elasticity of cancer cells with and without serum

Previous studies have reported that chromatin states modulate nuclear elasticity and that serum influences these chromatin states^59–61^. To investigate the effects of serum depletion on nuclear elasticity, we measured the elasticity of the intact nuclear surface in human lung cancer cells (PC9, harboring the EGFR Δexon19)^62^. Fig. 2a illustrates the NE-AFM method used for these measurements, where 256 force curves were taken at 16 × 16 arrayed-*xy* positions over a 1 × 1 μm^2^ area around the nucleus center. The set point of the force curve measurements (0.6 - 0.8 nN) was determined such that the tip approach stops at the nuclear surface, minimizing the risk of tip or cell damage. Thus, a two-dimensional map of the lowest tip heights corresponds to a height map of the nuclear surface, as shown in Fig. 2b.

**Fig. 2.**
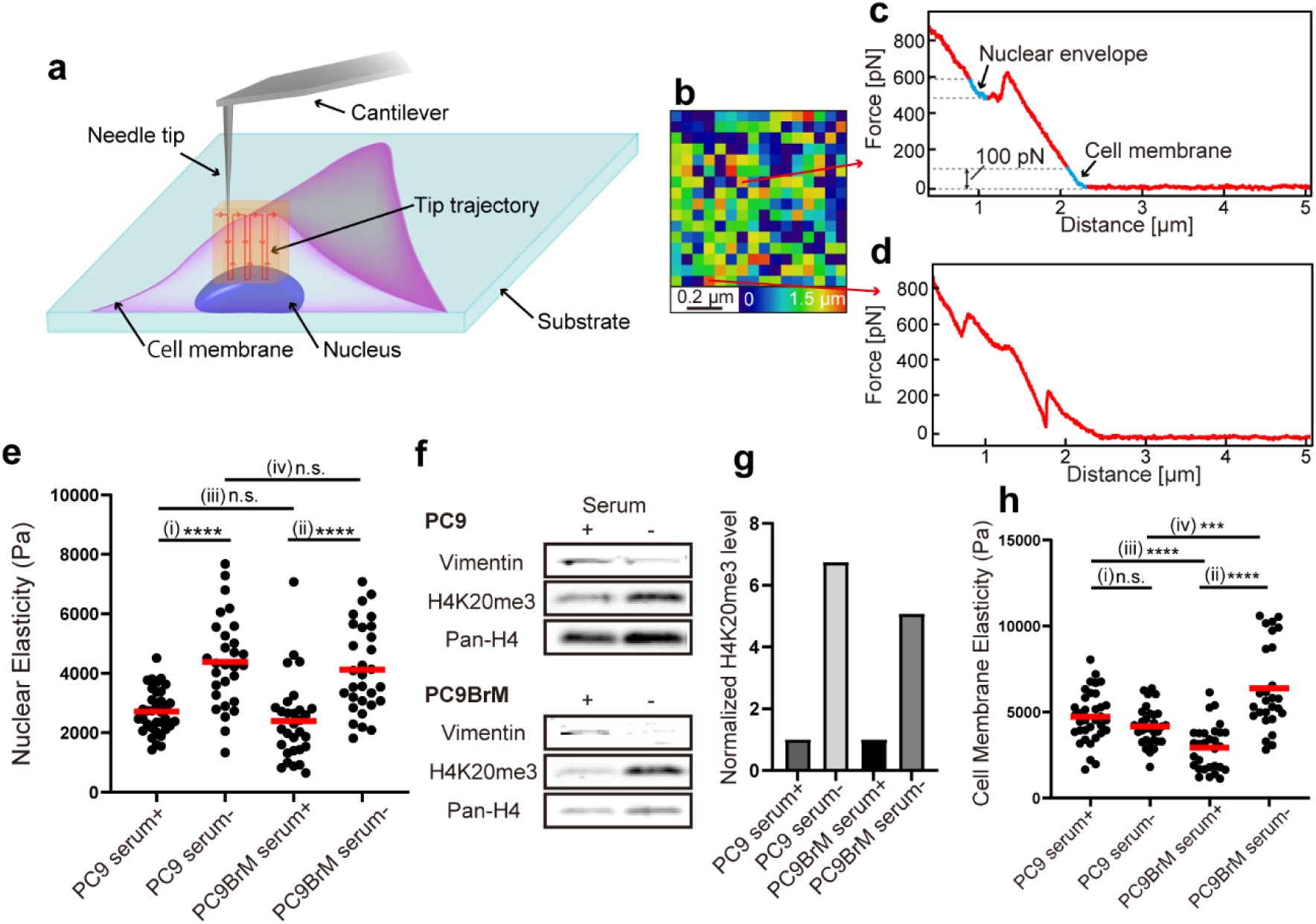
Quantitative analysis of nuclear elasticity by NE-AFM. (a) Schematic illustration of NE-AFM measurement on a nuclear surface. (b) Height map of the nucleus. (c) An example of a “good” force curve allowing unambiguous determination of the cell and nuclear CPs. (d) An example of a “bad” force curve is where identification of the cell or nuclear CP is difficult. (e) The distribution of nuclear elasticity of PC9 and PC9-BrM cells in normal (serum+) and serum-depletion (serum-) conditions. (f) Representative expression levels of vimentin, H4K20me3, and pan-H4 in PC9 and PC9-BrM with and without serum. (g) Bar graph of quantified levels of H4K20me3 normalized to pan-H4 from (g) by densitometry. Each bar was normalized to the expression level under serum+ conditions in PC9 and PC9BrM, respectively. (h) Distribution of cell membrane elasticity in PC9 and PC9-BrM cells under serum+ and serum-conditions.

Among the obtained 256 curves, some displayed a clear peak corresponding to cell membrane penetration, followed by a plateau and a sharp increase indicative of nuclear indentation (Fig. 2c). For such “good” curves, we could reliably identify the cell and nuclear CPs and estimate *E*_Y_. However, some “bad” curves show multiple small peaks due to the interaction with other intracellular components, making it difficult to reliably identify CPs and estimate *E*_Y_ (Fig. 2d). Therefore, we manually selected 20 good curves from the 256 curves and estimated the cell and nucleus *E*_Y_ by fitting the Hertz-Sneddon model to the indentation profiles (blue lines in Fig. 2c) using a fixed force range of 100 pN. The indentation depth of the nucleus ranged from 150 to 250 nm. As this range is comparable to the thickness of the nuclear envelope and associated chromatin structures^63–65^, the measurements should largely reflect their elasticity.

We measured the nuclear elasticity of living PC9 cells in culture media with and without serum. After two days of serum-free culture, nuclear elasticity significantly increased compared to conditions with serum (Fig. 2e(i) and Supplementary Table 1, *p* < 0.0001; with serum: 2,714 ± 126 Pa (average ± standard error of the mean, *N* = 35); without serum: 4,384 ± 284 Pa (*N* = 29)). We hypothesized that this increase in nuclear elasticity was due to changes in chromatin compaction. To test this, we measured levels of H4K20me3, a marker that increases with chromatin compaction, via immunoblotting^33,60^. H4K20me3 levels significantly increased under serum-depleted conditions compared to those before the serum depletion (Figs. 2f and g), indicating that serum depletion enhances chromatin compaction, leading to increased nuclear elasticity.

We performed similar experiments on brain-metastatic cells (PC9-BrM), which were established after four cycles of intra-cardiac injection and cancer cell collection^66^. Through these brain metastasis cycles, PC9 lung cancer cells likely acquired the ability to invade the blood-brain barrier (BBB), a network of endothelial cells with continuous tight junctions that represents the rate-limiting step in the development of brain metasiasis^67,68^. To go through the tight channels, these brain-metastatic cells may acquire reduced nuclear elasticity, as previously reported for other cancer cells^69^. Consistent with PC9 cells, serum depletion increased nuclear elasticity in PC9-BrM cells (Fig. 2e(ii), *p* < 0.0001; with serum: 2,394 ± 249 (*N* = 30); without serum: 4,122 ± 269 (*N* = 30)). H4K20me3 levels also increased under serum-depleted conditions (Figs. 2f and g). Unexpectedly, no significant differences in nuclear elasticity were observed between PC9 and PC9-BrM cells, regardless of serum presence (Fig. 2e(iii), *p* = 0.47; Fig. 2e(iv), *p* = 0.06).

Fig. 2h presents the results of cell membrane elasticity. In PC9 cells, cell membrane elasticity did not differ significantly between the presence and absence of serum (Fig. 2h(i), *p* = 0.08; with serum: 4,722 ± 250 (*N* = 35); without serum: 4,166 ± 198 (*N* = 31)). However, in PC9-BrM cells, cell membrane elasticity was significantly higher under serum-depleted conditions than under normal serum-treated conditions (Fig. 2h(ii), *p* < 0.001; with serum: 2,929 ± 241 (*N* = 30); without serum: 6,371 ± 477 (*N* = 27)). Increased expression levels of vimentin have been reported to enhance cell elasticity^70,71^. However, under serum-depleted conditions, vimentin expression was reduced (Fig. 2f), indicating that vimentin is not the main factor in increased cell membrane elasticity in PC9-BrM cells. When comparing PC9 and PC9-BrM cells, PC9-BrM cells exhibited lower cell membrane elasticity than PC9 cells in the presence of serum but higher elasticity under serum-free conditions (Fig. 2h(iii), *p* < 0.0001 (with serum); Fig. 2h(iv), *p* < 0.001 (without serum)). This means that PC9-BrM cells respond differently to serum compared with PC9, which may partially explain the discrepancy regarding previous cell elasticity experiments^7–9,72,73^.

### EMT induction using TGF-β and expression of lamins and histone modifications

Malignant transformation of cancer often requires epithelial-mesenchymal transition (EMT) induction^38–40,74^. To explore the relationship between malignant transformation and nuclear elasticity transitions, we examined whether EMT induction alters nuclear elasticity. EMT was induced by adding transforming growth factor (TGF)-β to a serum-free medium^39,40^. EMT induction was confirmed by the upregulation of vimentin and N-cadherin (Fig. 3e). Nuclear elasticity significantly decreased following EMT induction compared to control (serum-free) cells (Fig. 3a, *p* = 0.0018; 3,201 ± 197 Pa (*N* = 30)). A representative elasticity map of the nuclear surface in control and TGF-β treated cells on a 1 × 1 μm^2^ area is shown in Fig. 3b. In contrast, cell membrane elasticity remained unchanged (Fig. 3c, *p* = 0.513; 4,097 ± 302 Pa (*N* = 30); the elasticity map is shown in Fig. 3d).

**Fig. 3.**
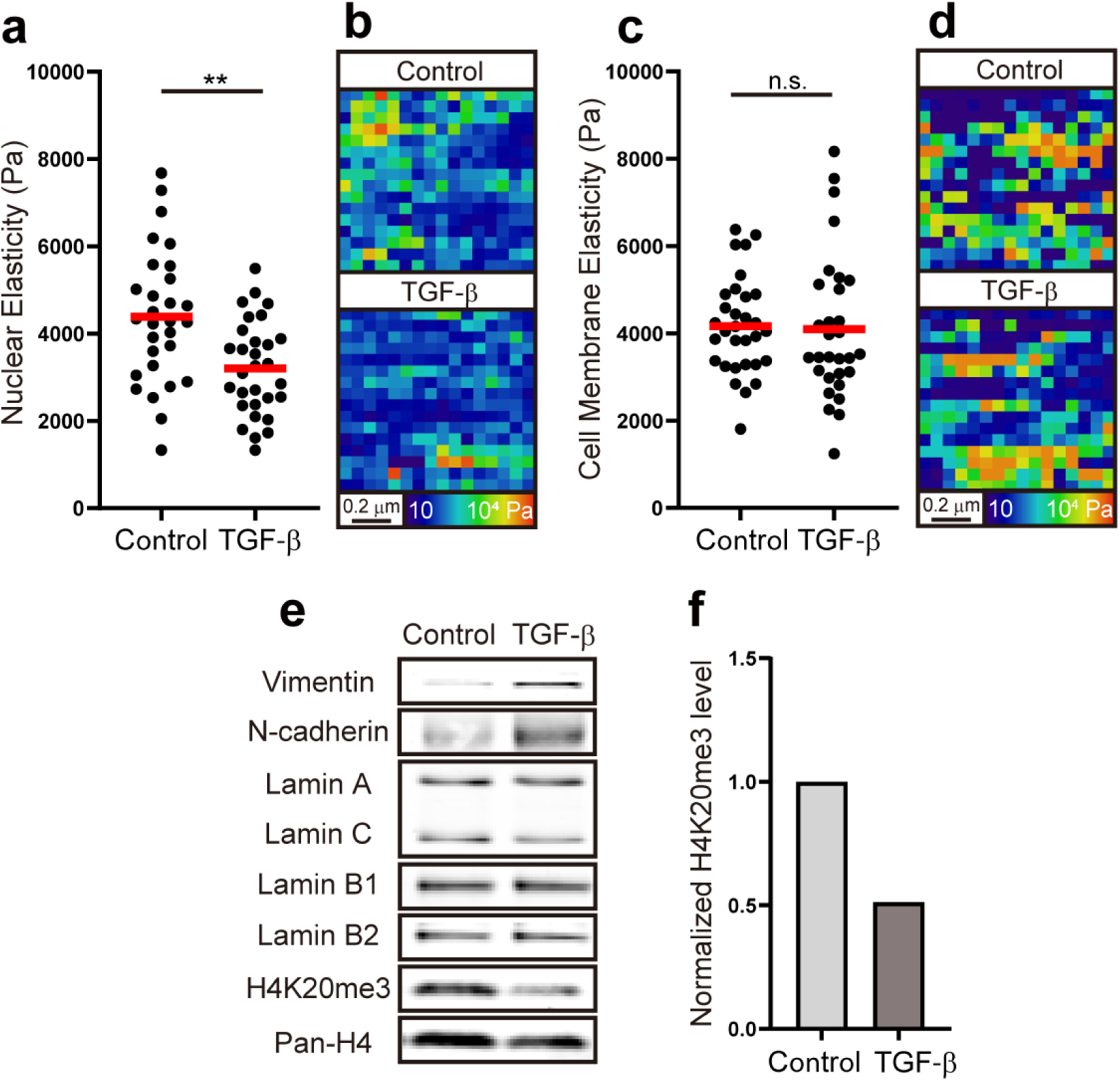
Measurement of the effect of EMT induction by TGF-β on nuclear and cell elasticity, as well as on lamins and histone modifications. (a) Nuclear elasticity of control (serum-free) and TGF-β treated cells. (b) Representative elasticity map of control and TGF-β treated cell nuclear surface. (c) Cell membrane elasticity. (d) Representative elasticity map of the cell membrane. (e) Expression levels of EMT markers (vimentin and N-cadherin), lamins, H4K20me3, and pan-H4. (f) Densitometrically quantified graph of H4K20me3 normalized to pan-H4 from (e).

Previous studies have shown that the NL plays a crucial role in regulating nuclear elasticity^22,59,75–77^. To determine whether the expression levels of lamins (lamin A/C, B1, and B2) are affected by TGF-β treatment, we measured their expression levels by immunoblotting. As shown in Fig. 3e, the expression levels of lamin A/C, B1, and B2 remained unchanged between control and TGF-β treated cells. This suggests that lamins are not the primary factor regulating nuclear elasticity under these conditions. In contrast, H4K20me3 levels decreased in PC9 cells after EMT induction (Figs. 3e and f). These results support the hypothesis that chromatin compaction contributes to alterations in nuclear elasticity.

### Measurement of nuclear volume, nuclear deformation, and cell adhesion area

We investigated the effects of serum and TGF-β on nuclear volume, nuclear deformation, and cell spreading to assess morphological changes in the cells. Fig. 4a shows fluorescence images of nuclei under serum treatment, serum depletion, and TGF-β treatment, while Figs. 4b and c present the distribution of nuclear volume and circularity under each condition. The nuclear volume of serum-depleted PC9 cells was significantly smaller than that of serum-treated cells. This reduction in nuclear volume may lead to increased nuclear elasticity by raising the concentration of structural components in the nuclear envelope. In contrast, no significant differences in nuclear volume were observed between serum-depleted cells and TGF-β-treated cells. Circularity, calculated by 4π × (area/perimeter^2^), quantifies how closely the shape resembles a perfect circle and serves as an indicator of nuclear deformation^78^. The results showed no significant differences in circularity under these three conditions. Fig. 4d presents representative fluorescence images of the cells under each condition, while Fig. 4e shows the distribution of cell adhesion area (the 2D projected area of a cell). The cell adhesion area was significantly reduced after serum depletion and increased following TGF-β treatment, likely due to enhanced lamellipodia spreading in response to serum and TGF-β. Overall, the presence or absence of serum and TGF-β did not significantly affect the shape of the nuclei or cells, nor did these factors influence the stiffness of the nuclear membrane.

**Fig. 4.**
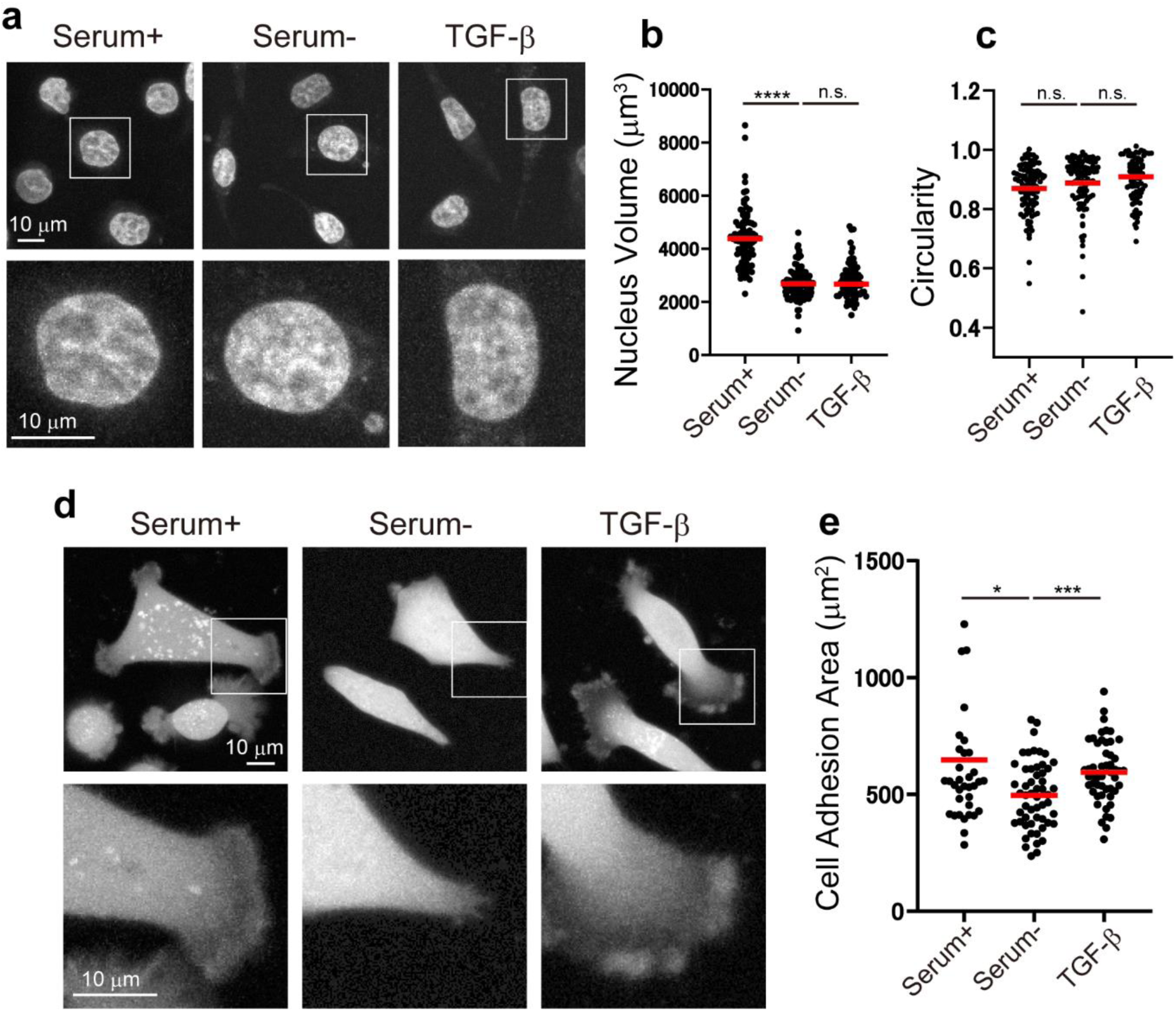
Quantification of nuclear volume, nuclear deformation, and cell adhesion area under serum (Serum+), serum depletion (Serum-), and TGF-β treatment. (a) Representative fluorescence images of nuclei for each treatment condition are shown as maximum-intensity projections. The images in the bottom row are magnified views of the area indicated by white boxes. (b) Distribution of nuclear volume. (c) Distribution of the nuclear circularity. (d) Representative fluorescence images of the cell under serum-containing medium, serum depletion, and TGF-β treatment. (e) Distribution of cell adhesion area.

## Discussion

In this study, we employed NE-AFM, which we recently developed to measure the nanoscale mechanical properties of nuclear elasticity in intact nuclei within living cancer cells. Our findings demonstrate that nuclear elasticity decreases when cells cultured without serum are exposed to serum or TGF-β. Notably, we found no significant changes in the expression levels of lamins A/C, B1, and B2 under these conditions. However, we observe a significant increase in H4K20me3 levels. These findings strongly suggest that chromatin compaction states influence nuclear elasticity. Exposure to serum or TGF-β reduces chromatin compaction, resulting in nuclear softening.

NE-AFM successfully mapped the elasticity distribution on the nuclear surface in living cells. The resulting elasticity map reveals highly heterogeneous distributions of nuclear elasticity (Figs. 1k, 3b, and d). The NL has a meshwork structure with a pore size of 1 – 1.5 μm in diameter^46^, and nuclear pore complexes (NPCs) are localized within these pores in the meshes^79^. Heterochromatin tethered to the NL forms lamina-associated domains (LADs)^80^, while regions near NPCs are enriched in euchromatin^48^. It is reasonable to speculate that these structural features contribute to the heterogeneous distributions of nuclear elasticity. Further experiments are required to elucidate the precise relationship between nuclear elasticity and these underlying structures. Our method offers enhanced precision for such investigations.

To date, no reports have examined the effect of serum and TGF-β on nuclear elasticity. In this study, we demonstrated that nuclear elasticity increases in the serum-depleted condition compared to serum-containing conditions in both parental and brain-metastatic PC9 cells (PC9 and PC9-BrM, respectively), and H4K20me3 levels increased in serum-depleted cells. A previous study reported that serum depletion increases H4K20me3 levels, supporting our model in which serum depletion increases H4K20me3 levels, leading to increased heterochromatin and, consequently, nuclear elasticity^61^. We also investigated the impact of TGF-β on nuclear elasticity in serum-free conditions and found that the TGF-β treatment reduces nuclear elasticity in both PC9 and PC9-BrM cells compared to control cells. Previous studies have reported that TGF-β decreases H4K20me3 levels^81^ and triggers a widespread increase in chromatin accessibility^82^, consistent with our findings. These results strongly suggest that TGF-β promotes global chromatin relaxation, leading to a decrease in nuclear elasticity.

When comparing nuclear elasticity between parental and brain-metastatic cells, we found that brain-metastatic cells (PC9-BrM) exhibit similar nuclear elasticity to parental PC9 cells. This result may seem unexpected, as EMT induction has been shown to decrease nuclear elasticity, and traversing the BBB is mediated by EMT^83^. One possible explanation is that PC9-BrM cells may have undergone EMT temporarily to traverse the BBB but gradually lost their metastatic properties during the subsequent development of brain metastases, cell collection, and repeated passaging in plastic culture dishes^84^. By the time nuclear elasticity was measured, PC9-BrM cells may have already lost their metastatic characteristics. On the other hand, cancer cells that underwent TGF-β-induced EMT directly are more likely to reflect EMT-associated properties.

The cell membrane elasticity of PC9-BrM cells exhibited opposite trends depending on the presence or absence of serum (Fig. 2h); the cell membrane elasticity of PC9-BrM cells was significantly lower than that of PC9 in the presence of serum, whereas it was higher in its absence. This observation may help explain conflicting results on cancer cell elasticity measurements. For instance, one study reported that the elasticity of cervical cancer cells was higher than normal cells when both were cultured in keratinocyte serum-free medium (KSFM)^73^, while another found that cervical cancer cells cultured in a medium containing 10% fetal bovine serum were softer than normal cells cultured in KSFM^16^. These findings suggest that differences in culture medium, particularly the presence or absence of serum, can significantly influence cell elasticity measurements. Therefore, we strongly recommend conducting all cell and nuclear elasticity measurements under controlled serum conditions.

The transition in the nuclear elasticity has been investigated as a potential biomarker for malignant cancer^14,15^. Accurate measurement of nuclear elasticity could significantly enhance clinical assessment. The method developed in this study could become a fundamental technique in cancer diagnostics and has broader applications for measuring the mechanical properties of other intracellular structures. While the mechanical properties of structures within living cells remain largely unexplored, our technique can be applied to assess the elasticity and adhesion properties of other organelle structures, such as mitochondrial membranes or focal adhesions, in both healthy and diseased cells^85,86^. Understanding these mechanical properties will provide deeper insights into nanoscale biology and the mechanisms underlying diseases associated with cellular dysfunction^87^.

## Methods

### Cell sample preparation

HeLa cells were obtained from the Japanese Collection of Research Bioresources (JCRB) Cell Bank (JCRB9004). PC9-Luc-EGFP and PC9-Luc-EGFP-BrM4 (brain-metastatic cells, referred to as PC9 and PC9-BrM, respectively, throughout the article) were established in a previous study^66^. All cell lines were maintained in Dulbecco’s Modified Eagle’s Medium (DMEM, Gibco) supplemented with 10% fetal bovine serum (Biosera) and 1% penicillin/streptomycin (Fujifilm Wako).

### NE-AFM

PC9 and PC9-BrM cells were cultured on 35-mm plastic dishes in the DMEM (Fujifilm Wako) containing 10% serum, no serum, or no serum and 5 ng/ml TGF-β1 (Peprotech) for two days. Before observation, the culture medium was replaced with Leibovitz’s L-15 medium (Thermo Fisher Scientific) supplemented with 1% penicillin/streptomycin and containing either 10% serum, no serum, or no serum with 5 ng/ml TGF-β1. All experiments were conducted within a few hours after the medium change. The nuclear and cell membrane elasticities were measured using NE-AFM methods^55–57^. Nanoneedle probes were fabricated using electron beam deposition with a focused ion beam system (Helios G4 CX Dual Beam, Thermo Fisher Scientific) on tip-truncated cantilevers (Olympus, BLAC40TS-C2, spring constant approximately 0.1 N/m). We used a JPK NanoWizard 4 BioAFM (Bruker) equipped with an inverted fluorescence microscope (Eclipse Ti2, Nikon). The temperature was maintained at 37 °C using a dish heater (Bruker). Measurements were conducted in QI mode with the following parameters: 16 × 16 pixels, 5.5 – 6.5 μm Z-length, 1.5 nN setpoint for whole cell or 0.6 – 0.8 nN for nuclear elasticity, and 10 μm/s tip speed. 10 cells were measured per sample, and 20 force curves were selected out of 256 curves for each cell for *E*_Y_ estimation. *E*_Y_ was calculated by fitting the force carves within a fixed 0.1 nN range using the following equation with custom software^25^. *F*, *Rc*, ν, and δ indicate force, the probe’s radius, Poisson ratio, and indentation depth, respectively^57^.

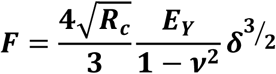

### Immunoblotting

Cultured cells on 10-cm dishes were washed three times with ice-cold PBS and lysed in 150 μl of RIPA or Laemmli buffer supplemented with protease inhibitor and phosphatase inhibitor cocktail (cOmplete and PhosSTOP, Roche). After lysis, samples were sonicated for two minutes, freeze-thaw once, and heated at 95°C for five minutes. Protein concentration was measured using a BCA Protein Assay Kit (BioDynamics Laboratory), and samples were adjusted to equal concentrations. Samples were mixed with 0.005% bromophenol blue (Nacalai) and 125 mM DTT (Fujifilm Wako), then denatured at 95°C for five minutes. Proteins (3-20 μg) were separated on 5-20% polyacrylamide gels (SuperSep Ace, Fujifilm Wako) with a marker (Precision Plus Protein, Bio-Rad) and transferred onto polyvinylidene fluoride (PVDF) or nitrocellulose membrane (Bio-Rad) using a Trans-Blot Turbo semi-dry transfer system (Bio-Rad). Membranes were blocked with Intercept Blocking Buffer (TBS; LI-COR) for one hour at room temperature, incubated overnight at 4°C with primary antibodies, and then incubated with secondary antibodies for one hour at room temperature. Antibodies used for immunoblotting are listed in Supplemental Table 2. Fluorescence signals were detected using an Odyssey CLx imaging system (LI-COR). Total proteins were visualized with Ponceau S Staining Solution (Supplementary Fig. 1, Beacle), and images were analyzed using Image Studio Lite (LI-COR).

### Measurement of cell area and nucleus volume

Cells are cultured on a glass-bottom dish (ibidi). For measuring nuclear volume and nuclear deformation, the nuclei were stained with 1 μM SiR-DNA (Cytoskeleton). Fluorescence images were acquired in three dimensions using a confocal microscope (Expert Line, Abberior Instruments) with a resolution of 200 × 200 × 200 nm per pixel, using a 640 nm excitation and a 10 μs exposure time. Image analysis was performed using a custom MATLAB script (R2024a, Simulink). For measuring cell adhesion area, the cytosol was stained with 1 μg/ml Calcein-AM (Dojindo), and the cell membrane was stained with PlasMem Bright Green (1:1000 dilution, Dojindo). Scatter plots were generated using Prism 9 (GraphPad Software).

### Statistical analyses

Welch’s t-tests were used for single comparisons, and Games-Howell tests were used for multiple comparisons, both performed using EZR software (Saitama Medical Center, Jichi Medical University)^88^. The symbols n.s., *, **, ***, and **** indicate p-values as follows: n.s. (non-significant, *p* > 0.05), * (*p* < 0.05), ** (*p* < 0.01), *** (*p* < 0.001), and **** (*p* < 0.0001).

## Acknowledgments

This work was supported by the World Premier International Research Center Initiative (WPI), MEXT, Japan, by JSPS KAKENHI under grant numbers 20H00345, 21H05251, 22H01954, and 23K05763. Additional support was provided by Platforms for Advanced Technologies and Research Resources “Advanced Bioimaging Support” (JSPS KAKENHI Grant Number JP22H04926). Ichikawa was supported by the Mitani Foundation for Research and Development, Takeda Science Foundation, Shimadzu Science Foundation, and Nakatani Foundation. We thank Kiminori Toyooka and Mayuko Sato for the preliminary analysis.

## Author Contributions

1. T. I., Y. K., K. S., T. S., and T. F. designed the experiments. K. I. and E. H. prepared the cell lines. T. I. and K. S. performed the AFM experiment and analysis under the supervision of T. F.. Analysis software was developed by N. M. and T. M. under the supervision of K. M. and T. F.. A. K. and Y. K. performed immunoblotting under the supervision of K. I., E. H. and T. S.. H. K. produced mouse monoclonal anti-histone modification-specific antibodies. T. Y. did a preliminary analysis of the protein expression of the cell under the supervision of R. H.. T. I., Y. K. and T. S. wrote the manuscript. All authors read and approved the final manuscript.

## Additional information

Supplementary information: The online version contains supplementary material available at https://doi.org/.

## Preprints

This manuscript was posted on a preprint: https://doi.org/10.1101/2024.12.13.626302.

## Data availability

**Correspondence** and requests for materials should be addressed to Takehiko Ichikawa or Takeshi Fukuma.

**Peer review information** Nature Communications thanks Blaine Bartholomew and the other anonymous reviewer(s) for their contribution to the peer review of this work. A peer review file is available.

**Reprints and permissions information** is available at http://www.nature.com/reprints

**Publisher’s note:** Springer Nature remains neutral with regard to jurisdictional claims in published maps and institutional affiliations.

**Open Access**

## References

1 Sigrist, R. M. S., Liau, J., Kaffas, A. E., Chammas, M. C. & Willmann, J. K. Ultrasound Elastography: Review of Techniques and Clinical Applications. Theranostics 7, 1303–1329 (2017). 10.7150/thno.18650

2 Wells, P. N. & Liang, H. D. Medical ultrasound: imaging of soft tissue strain and elasticity. J R Soc Interface 8, 1521–1549 (2011). 10.1098/rsif.2011.0054

3 Fass, L. Imaging and cancer: a review. Mol Oncol 2, 115–152 (2008). 10.1016/j.molonc.2008.04.001

4 Levental, K. R. et al. Matrix crosslinking forces tumor progression by enhancing integrin signaling. Cell 139, 891–906 (2009). 10.1016/j.cell.2009.10.027

5 Yuan, Z. et al. Extracellular matrix remodeling in tumor progression and immune escape: from mechanisms to treatments. Mol Cancer 22, 48 (2023). 10.1186/s12943-023-01744-8

6 Elgundi, Z. et al. Cancer Metastasis: The Role of the Extracellular Matrix and the Heparan Sulfate Proteoglycan Perlecan. Front Oncol 9, 1482 (2019). 10.3389/fonc.2019.01482

7 Kashani, A. S. & Packirisamy, M. Cancer cells optimize elasticity for efficient migration. R Soc Open Sci 7, 200747 (2020). 10.1098/rsos.200747

8 Denais, C. & Lammerding, J. Nuclear mechanics in cancer. Adv Exp Med Biol 773, 435–470 (2014). 10.1007/978-1-4899-8032-8_20

9 Lekka, M. et al. Elasticity of normal and cancerous human bladder cells studied by scanning force microscopy. Eur Biophys J 28, 312–316 (1999). 10.1007/s002490050213

10 Liu, Z. et al. Cancer cells display increased migration and deformability in pace with metastatic progression. FASEB J 34, 9307–9315 (2020). 10.1096/fj.202000101RR

11 Dongre, A. & Weinberg, R. A. New insights into the mechanisms of epithelial-mesenchymal transition and implications for cancer. Nat Rev Mol Cell Biol 20, 69–84 (2019). 10.1038/s41580-018-0080-4

12 Leggett, S. E., Hruska, A. M., Guo, M. & Wong, I. Y. The epithelial-mesenchymal transition and the cytoskeleton in bioengineered systems. Cell Commun Signal 19, 32 (2021). 10.1186/s12964-021-00713-2

13 Leggett, S. E. et al. Morphological single cell profiling of the epithelial-mesenchymal transition. Integr Biol (Camb) 8, 1133–1144 (2016). 10.1039/c6ib00139d

14 Liu, S. et al. Mechanotherapy in oncology: Targeting nuclear mechanics and mechanotransduction. Adv Drug Deliv Rev 194, 114722 (2023). 10.1016/j.addr.2023.114722

15 Xu, W. et al. Cell stiffness is a biomarker of the metastatic potential of ovarian cancer cells. PLoS One 7, e46609 (2012). 10.1371/journal.pone.0046609

16 Zhao, X., Zhong, Y., Ye, T., Wang, D. & Mao, B. Discrimination Between Cervical Cancer Cells and Normal Cervical Cells Based on Longitudinal Elasticity Using Atomic Force Microscopy. Nanoscale Res Lett 10, 482 (2015). 10.1186/s11671-015-1174-y

17 Li, Q. S., Lee, G. Y., Ong, C. N. & Lim, C. T. AFM indentation study of breast cancer cells. Biochem Biophys Res Commun 374, 609–613 (2008). 10.1016/j.bbrc.2008.07.078

18 Rianna, C. & Radmacher, M. Comparison of viscoelastic properties of cancer and normal thyroid cells on different stiffness substrates. Eur Biophys J 46, 309–324 (2017). 10.1007/s00249-016-1168-4

19 Wang, S. Q. et al. Membrane Deformability and Membrane Tension of Single Isolated Mitochondria. Cell Mol Bioeng 1, 67–74 (2008). 10.1007/s12195-008-0002-1

20 Fischer, T., Hayn, A. & Mierke, C. T. Effect of Nuclear Stiffness on Cell Mechanics and Migration of Human Breast Cancer Cells. Front Cell Dev Biol 8, 393 (2020). 10.3389/fcell.2020.00393

21 Krause, M., Te Riet, J. & Wolf, K. Probing the compressibility of tumor cell nuclei by combined atomic force-confocal microscopy. Phys Biol 10, 065002 (2013). 10.1088/1478-3975/10/6/065002

22 Swift, J. et al. Nuclear lamin-A scales with tissue stiffness and enhances matrix-directed differentiation. Science 341, 1240104 (2013). 10.1126/science.1240104

23 Hobson, C. M. et al. Correlating nuclear morphology and external force with combined atomic force microscopy and light sheet imaging separates roles of chromatin and lamin A/C in nuclear mechanics. Mol Biol Cell 31, 1788–1801 (2020). 10.1091/mbc.E20-01-0073

24 Lanzicher, T. et al. AFM single-cell force spectroscopy links altered nuclear and cytoskeletal mechanics to defective cell adhesion in cardiac myocytes with a nuclear lamin mutation. Nucleus-Phila 6, 394–407 (2015). 10.1080/19491034.2015.1084453

25 Vinckier, A. & Semenza, G. Measuring elasticity of biological materials by atomic force microscopy. FEBS Lett 430, 12–16 (1998). 10.1016/s0014-5793(98)00592-4

26 Wang, K., Qin, Y. & Chen, Y. In situ AFM detection of the stiffness of the in situ exposed cell nucleus. Biochim Biophys Acta Mol Cell Res 1868, 118985 (2021). 10.1016/j.bbamcr.2021.118985

27 Beicker, K., O’Brien, E. T., 3rd, Falvo, M. R. & Superfine, R. Vertical Light Sheet Enhanced Side-View Imaging for AFM Cell Mechanics Studies. Sci Rep 8, 1504 (2018). 10.1038/s41598-018-19791-3

28 Wei, F., Lan, F., Liu, B., Liu, L. Q. & Li, G. Y. Poroelasticity of cell nuclei revealed through atomic force microscopy characterization. Appl Phys Lett 109 (2016). Artn 213701 10.1063/1.4968191

29 Zhou, G. Q., Zhang, B. K., Tang, G. L., Yu, X. F. & Galluzzi, M. Cells nanomechanics by atomic force microscopy: focus on interactions at nanoscale. Adv Phys-X 6 (2021). Artn 1866668 10.1080/23746149.2020.1866668

30 Ujihara, Y., Ono, D., Ito, M., Sugita, S. & Nakamura, M. Nuclear deformability of cancer cells with different metastatic potential. Journal of Biorheology 37, 56–63 (2023).

31 Rowat, A. C., Lammerding, J. & Ipsen, J. H. Mechanical properties of the cell nucleus and the effect of emerin deficiency. Biophys J 91, 4649–4664 (2006). 10.1529/biophysj.106.086454

32 Dahl, K. N., Kahn, S. M., Wilson, K. L. & Discher, D. E. The nuclear envelope lamina network has elasticity and a compressibility limit suggestive of a molecular shock absorber. J Cell Sci 117, 4779–4786 (2004). 10.1242/jcs.01357

33 Pajerowski, J. D., Dahl, K. N., Zhong, F. L., Sammak, P. J. & Discher, D. E. Physical plasticity of the nucleus in stem cell differentiation. Proc Natl Acad Sci U S A 104, 15619–15624 (2007). 10.1073/pnas.0702576104

34 Tang, W. et al. Indentation induces instantaneous nuclear stiffening and unfolding of nuclear envelope wrinkles. Proc Natl Acad Sci U S A 120, e2307356120 (2023). 10.1073/pnas.2307356120

35 Mazumder, A., Roopa, T., Basu, A., Mahadevan, L. & Shivashankar, G. V. Dynamics of chromatin decondensation reveals the structural integrity of a mechanically prestressed nucleus. Biophys J 95, 3028–3035 (2008). 10.1529/biophysj.108.132274

36 Gilbert, N. et al. Chromatin architecture of the human genome: gene-rich domains are enriched in open chromatin fibers. Cell 118, 555–566 (2004). 10.1016/j.cell.2004.08.011

37 Allshire, R. C. & Madhani, H. D. Ten principles of heterochromatin formation and function. Nat Rev Mol Cell Biol 19, 229–244 (2018). 10.1038/nrm.2017.119

38 Tian, H. et al. PHF14 enhances DNA methylation of SMAD7 gene to promote TGF-beta-driven lung adenocarcinoma metastasis. Cell Discov 9, 41 (2023). 10.1038/s41421-023-00528-0

39 Zhang, H. J. et al. Transforming growth factor-beta1 promotes lung adenocarcinoma invasion and metastasis by epithelial-to-mesenchymal transition. Mol Cell Biochem 355, 309–314 (2011). 10.1007/s11010-011-0869-3

40 Zhang, N. et al. Decitabine reverses TGF-beta1-induced epithelial-mesenchymal transition in non-small-cell lung cancer by regulating miR-200/ZEB axis. Drug Des Devel Ther 11, 969–983 (2017). 10.2147/DDDT.S129305

41 Williams, J. F. et al. The condensation of HP1alpha/Swi6 imparts nuclear stiffness. Cell Rep 43, 114373 (2024). 10.1016/j.celrep.2024.114373

42 Ungricht, R. & Kutay, U. Mechanisms and functions of nuclear envelope remodelling. Nat Rev Mol Cell Biol 18, 229–245 (2017). 10.1038/nrm.2016.153

43 Crisp, M. et al. Coupling of the nucleus and cytoplasm: role of the LINC complex. J Cell Biol 172, 41–53 (2006). 10.1083/jcb.200509124

44 Gerace, L. & Blobel, G. The nuclear envelope lamina is reversibly depolymerized during mitosis. Cell 19, 277–287 (1980). 10.1016/0092-8674(80)90409-2

45 Foisner, R. & Gerace, L. Integral membrane proteins of the nuclear envelope interact with lamins and chromosomes, and binding is modulated by mitotic phosphorylation. Cell 73, 1267–1279 (1993). 10.1016/0092-8674(93)90355-t

46 Shimi, T. et al. Structural organization of nuclear lamins A, C, B1, and B2 revealed by superresolution microscopy. Mol Biol Cell 26, 4075–4086 (2015). 10.1091/mbc.E15-07-0461

47 Shimi, T. et al. The A- and B-type nuclear lamin networks: microdomains involved in chromatin organization and transcription. Genes Dev 22, 3409–3421 (2008). 10.1101/gad.1735208

48 Schermelleh, L. et al. Subdiffraction multicolor imaging of the nuclear periphery with 3D structured illumination microscopy. Science 320, 1332–1336 (2008). 10.1126/science.1156947

49 Turgay, Y. et al. The molecular architecture of lamins in somatic cells. Nature 543, 261–264 (2017). 10.1038/nature21382

50 Srivastava, L. K., Ju, Z., Ghagre, A. & Ehrlicher, A. J. Spatial distribution of lamin A/C determines nuclear stiffness and stress-mediated deformation. J Cell Sci 134 (2021). 10.1242/jcs.248559

51 Obataya, I., Nakamura, C., Han, S., Nakamura, N. & Miyake, J. Mechanical sensing of the penetration of various nanoneedles into a living cell using atomic force microscopy. Biosens Bioelectron 20, 1652–1655 (2005). 10.1016/j.bios.2004.07.020

52 Obataya, I., Nakamura, C., Han, S., Nakamura, N. & Miyake, J. Nanoscale operation of a living cell using an atomic force microscope with a nanoneedle. Nano Lett 5, 27–30 (2005). 10.1021/nl0485399

53 Liu, H. et al. In situ mechanical characterization of the cell nucleus by atomic force microscopy. ACS Nano 8, 3821–3828 (2014). 10.1021/nn500553z

54 Wang, X. et al. Mechanical stability of the cell nucleus - roles played by the cytoskeleton in nuclear deformation and strain recovery. J Cell Sci 131 (2018). 10.1242/jcs.209627

55 Penedo, M. et al. Visualizing intracellular nanostructures of living cells by nanoendoscopy-AFM. Sci Adv 7, eabj4990 (2021). 10.1126/sciadv.abj4990

56 Penedo, M. et al. Cell penetration efficiency analysis of different atomic force microscopy nanoneedles into living cells. Sci Rep 11, 7756 (2021). 10.1038/s41598-021-87319-3

57 Ichikawa, T. et al. Protocol for live imaging of intracellular nanoscale structures using atomic force microscopy with nanoneedle probes. STAR Protoc 4, 102468 (2023). 10.1016/j.xpro.2023.102468

58 Sneddon, I. N. The relation between load and penetration in the axisymmetric Boussinesq problem for a punch of arbitrary profile. International journal of engineering science 3, 47–57 (1965).

59 Stephens, A. D., Banigan, E. J., Adam, S. A., Goldman, R. D. & Marko, J. F. Chromatin and lamin A determine two different mechanical response regimes of the cell nucleus. Mol Biol Cell 28, 1984–1996 (2017). 10.1091/mbc.E16-09-0653

60 Stephens, A. D. et al. Chromatin histone modifications and rigidity affect nuclear morphology independent of lamins. Mol Biol Cell 29, 220–233 (2018). https://doi.org:10.1091/mbc.E17-06-0410

61 Kourmouli, N. et al. Heterochromatin and tri-methylated lysine 20 of histone H4 in animals. J Cell Sci 117, 2491–2501 (2004). 10.1242/jcs.01238

62 Kinjo, M. et al. Thromboplastic and fibrinolytic activities of cultured human cancer cell lines. Br J Cancer 39, 15–23 (1979). 10.1038/bjc.1979.3

63 Erenpreisa, J. et al. Differential staining of peripheral nuclear chromatin with Acridine orange implies an A-form epichromatin conformation of the DNA. Nucleus-Phila 9, 171–181 (2018). 10.1080/19491034.2018.1431081

64 Li, Y. et al. Analysis of three-dimensional chromatin packing domains by chromatin scanning transmission electron microscopy (ChromSTEM). Sci Rep 12, 12198 (2022). 10.1038/s41598-022-16028-2

65 De Boni, U. Chromatin and nuclear envelope of freeze-fractured, neuronal interphase nuclei, resolved by scanning electron microscopy. Biol Cell 63, 1–8 (1988). 10.1111/j.1768-322x.1988.tb00735.x

66 Hirata, E. et al. The Brain Microenvironment Induces DNMT1 Suppression and Indolence of Metastatic Cancer Cells. iScience 23, 101480 (2020). 10.1016/j.isci.2020.101480

67 Abbott, N. J., Patabendige, A. A., Dolman, D. E., Yusof, S. R. & Begley, D. J. Structure and function of the blood-brain barrier. Neurobiol Dis 37, 13–25 (2010). 10.1016/j.nbd.2009.07.030

68 Xia, S. et al. Mesothelin promotes brain metastasis of non-small cell lung cancer by activating MET. J Exp Clin Cancer Res 43, 103 (2024). 10.1186/s13046-024-03015-w

69 Kwon, S., Yang, W., Moon, D. & Kim, K. S. Comparison of Cancer Cell Elasticity by Cell Type. J Cancer 11, 5403–5412 (2020). 10.7150/jca.45897

70 Patteson, A. E. et al. Vimentin protects cells against nuclear rupture and DNA damage during migration. J Cell Biol 218, 4079–4092 (2019). 10.1083/jcb.201902046

71 Pogoda, K. et al. Unique Role of Vimentin Networks in Compression Stiffening of Cells and Protection of Nuclei from Compressive Stress. Nano Lett 22, 4725–4732 (2022). 10.1021/acs.nanolett.2c00736

72 Zhang, G., Long, M., Wu, Z. Z. & Yu, W. Q. Mechanical properties of hepatocellular carcinoma cells. World J Gastroenterol 8, 243–246 (2002). 10.3748/wjg.v8.i2.243

73 Iyer, S., Gaikwad, R. M., Subba-Rao, V., Woodworth, C. D. & Sokolov, I. Atomic force microscopy detects differences in the surface brush of normal and cancerous cells. Nat Nanotechnol 4, 389–393 (2009). 10.1038/nnano.2009.77

74 Brabletz, T., Kalluri, R., Nieto, M. A. & Weinberg, R. A. EMT in cancer. Nat Rev Cancer **18**, 128–134 (2018). 10.1038/nrc.2017.118

75 Broers, J. L. et al. Decreased mechanical stiffness in LMNA-/- cells is caused by defective nucleo-cytoskeletal integrity: implications for the development of laminopathies. Hum Mol Genet 13, 2567–2580 (2004). 10.1093/hmg/ddh295

76 Lammerding, J. et al. Lamins A and C but not lamin B1 regulate nuclear mechanics. J Biol Chem 281, 25768–25780 (2006). 10.1074/jbc.M513511200

77 Ovsiannikova, N. L. et al. Lamin A as a Determinant of Mechanical Properties of the Cell Nucleus in Health and Disease. Biochemistry (Mosc) 86, 1288–1300 (2021). 10.1134/S0006297921100102

78 Kamikawa, Y. et al. Impact of cell cycle on repair of ruptured nuclear envelope and sensitivity to nuclear envelope stress in glioblastoma. Cell Death Discov 9, 233 (2023). 10.1038/s41420-023-01534-7

79 Shevelyov, Y. Y. Interactions of Chromatin with the Nuclear Lamina and Nuclear Pore Complexes. Int J Mol Sci 24 (2023). 10.3390/ijms242115771

80 Guelen, L. et al. Domain organization of human chromosomes revealed by mapping of nuclear lamina interactions. Nature 453, 948–951 (2008). 10.1038/nature06947

81 Lyu, G. et al. Addendum: TGF-beta signaling alters H4K20me3 status via miR-29 and contributes to cellular senescence and cardiac aging. Nat Commun 9, 4134 (2018). 10.1038/s41467-018-06710-3

82 Guerrero-Martinez, J. A., Ceballos-Chavez, M., Koehler, F., Peiro, S. & Reyes, J. C. TGFbeta promotes widespread enhancer chromatin opening and operates on genomic regulatory domains. Nat Commun 11, 6196 (2020). 10.1038/s41467-020-19877-5

83 Jeevan, D. S., Cooper, J. B., Braun, A., Murali, R. & Jhanwar-Uniyal, M. Molecular Pathways Mediating Metastases to the Brain via Epithelial-to-Mesenchymal Transition: Genes, Proteins, and Functional Analysis. Anticancer Res 36, 523–532 (2016).

84 Acharekar, A. et al. Substrate stiffness regulates the recurrent glioblastoma cell morphology and aggressiveness. Matrix Biol 115, 107–127 (2023). 10.1016/j.matbio.2022.12.002

85 Feng, Q. & Kornmann, B. Mechanical forces on cellular organelles. J Cell Sci 131 (2018). 10.1242/jcs.218479

86 Bartolak-Suki, E., Imsirovic, J., Nishibori, Y., Krishnan, R. & Suki, B. Regulation of Mitochondrial Structure and Dynamics by the Cytoskeleton and Mechanical Factors. Int J Mol Sci 18 (2017). 10.3390/ijms18081812

87 Meyers, M. A., Chen, P. Y., Lopez, M. I., Seki, Y. & Lin, A. Y. Biological materials: a materials science approach. J Mech Behav Biomed Mater 4, 626–657 (2011). 10.1016/j.jmbbm.2010.08.005

88 Kanda, Y. Investigation of the freely available easy-to-use software ’EZR’ for medical statistics. Bone Marrow Transplant 48, 452–458 (2013). 10.1038/bmt.2012.244

