## Supplementaly Information for "Probing nanomechanics by direct indentation using Nanoendoscopy-AFM reveals the nuclear elasticity transition in cancer cells"

*Correspondence

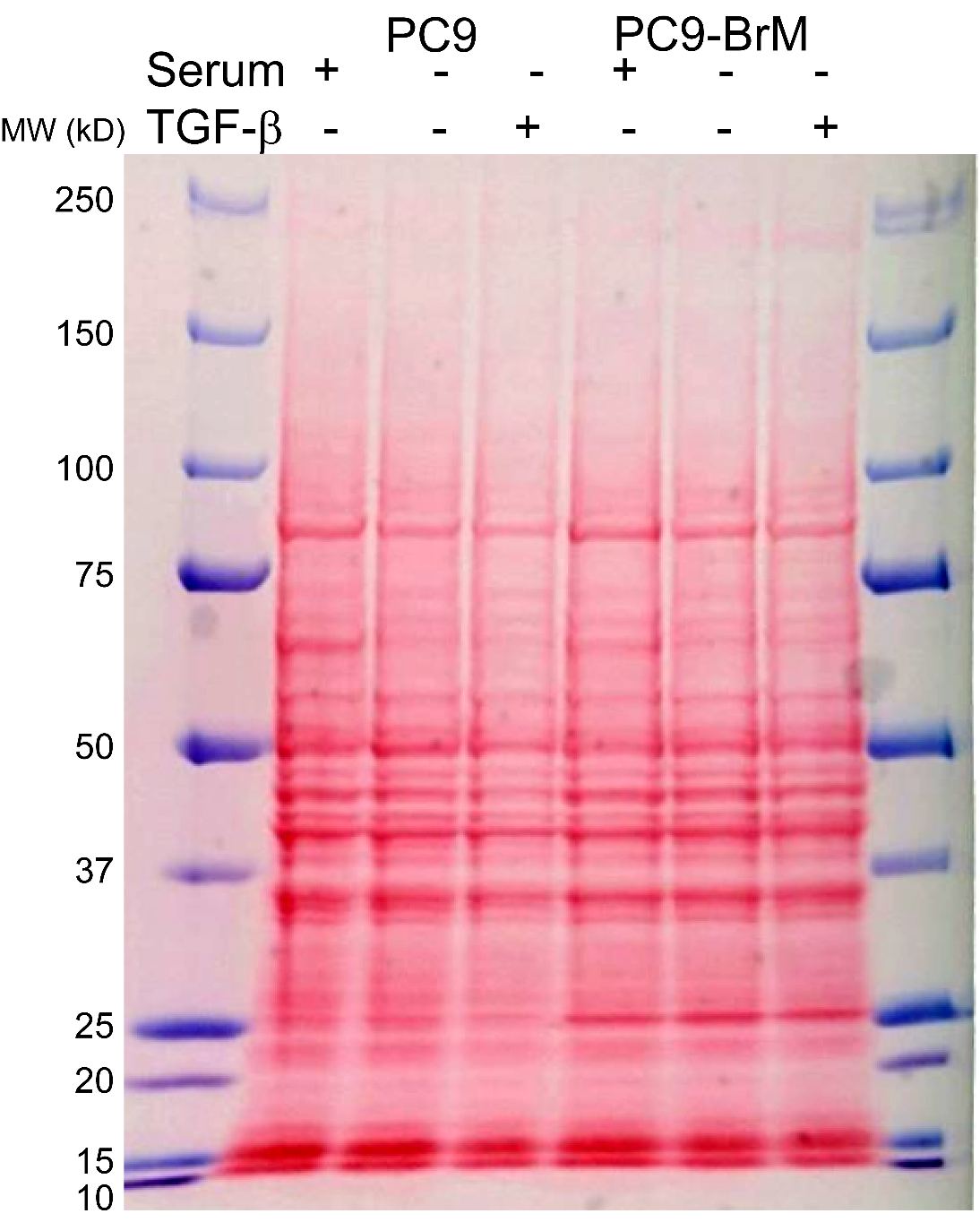

**Supplemental Fig. 1 |** Total proteins stained with Ponceau S related with Figs. 2f and 3e. MW: molecular weight.

| Pa ± SEM | Serum+ | Serum- | TGF-β |
| --- | --- | --- | --- |
| PC9 nucleus | 2714 ± 126 | 4384 ± 284 | 3201 ± 198 |
| PC9-BrM nucleus | 2394 ± 249 | 4122 ± 269 |  |
| PC9 cell membrane | 4722 ± 250 | 4166 ± 198 | 4097 ± 302 |
| PC9-BrM cell membrane | 2929 ± 241 | 6371 ± 477 |  |

**Supplemental Table 1 |** Young’s modulus of the nuclear surface and cell membrane under serum-present (Serum+), serum-absent (Serum-), and TGF-β-treated conditions.

| Primary antibody | | | | |
| --- | --- | --- | --- | --- |
| Target | Host | Clone/Name (Source) | RRID | Dilution rate |
| Vimentin | Rabbit | D21H3 (Cell Signaling) | AB_10695459 | 1:500 |
| N-cadherin | Mouse | 13A9 (Santa Cruz) | AB_781744 | 1:200 |
| Lamin A/C | Mouse | 3A6-4C11 (eBioscience) | AB_2802197 | 1:20,000 |
| Lamin B1 | Rabbit | 12987-1-AP (Proteintech) | AB_2136290 | 1:6,000 |
| Lamin B2 | Mouse | 8F6-E8-F12 (Biolegend) | AB_2832882 | 1;1,000 |
| H4K20me3 | Mouse | CMA423/27F10  (Hayashi-Takanaka et al. 2015)^1^ |  | 1:6,000 |
| Pan-H4 | Mouse | CMA400/9C5  (Hayashi-Takanaka et al. 2015)^1^ |  | 1:5,000 |
| Secondary antibody | | | | |
| Name (Source) | | | RRID | Dilution rate |
| IRDye 800CW Donkey anti-Mouse IgG (LI-COR) | | | AB_621847 | 1:20,000 |
| IRDye 680RD Donkey anti-Rabbit IgG (LI-COR) | | | AB_10954442 | 1:20,000 |

**Supplemental Table 2 |** Antibody list used for immunoblotting.

| Fig. 2e Nuclear elasticity | | | |
| --- | --- | --- | --- |
| Group | *N* |  | |
| PC9serum+ | 35 |  |  |
| PC9serum- | 29 |  |  |
| PC9BrMserum+ | 30 |  |  |
| PC9BrMserum- | 30 |  |  |
| Welch’s one-way ANOVA | *df* | *F* | *p* |
| Factor (Group) | 3 | 17.93 | 0.0000000011 |
| Residual | 120 |  |  |
| Games-Howell test | *df* | *t* | *p* |
| PC9serum+ vs. PC9serum- | 38.81 | 5.37 | 0.000023 |
| PC9BrMserum+ vs. PC9BrMserum- | 57.65 | 4.71 | 0.000093 |
| PC9serum+ vs. PC9BrMserum+ | 43.28 | 1.15 | 0.66 |
| PC9serum- vs. PC9BrMserum- | 56.71 | 0.67 | 0.91 |
| PC9serum+ vs. PC9BrMserum- | 41.35 | 4.74 | 0.00015 |
| PC9serum- vs. PC9BrMserum+ | 55.76 | 5.263 | 0.000014 |
| Fig. 2h Cell membrane elasticity | | | |
| Group | *N* |  | |
| PC9serum+ | 35 |  |  |
| PC9serum- | 31 |  |  |
| PC9BrMserum+ | 30 |  |  |
| PC9BrMserum- | 27 |  |  |
| Welch’s one-way ANOVA | *df* | *F* | *p* |
| Factor (Group) | 3 | 21.45 | 0.000000000035 |
| Residual | 119 |  |  |
| Games-Howell test | *df* | *t* | *p* |
| PC9serum+ vs. PC9serum- | 62.29 | 1.74 | 0.31 |
| PC9BrMserum+ vs. PC9BrMserum- | 38.73 | 6.44 | 0.00000077 |
| PC9serum+ vs. PC9BrMserum+ | 62.87 | 5.16 | 0.000016 |
| PC9serum- vs. PC9BrMserum- | 34.84 | 4.27 | 0.00079 |
| PC9serum+ vs. PC9BrMserum- | 39.92 | 3.06 | 0.020 |
| PC9serum- vs. PC9BrMserum+ | 56.46 | 3.96 | 0.0012 |

**Supplemental Table 3 |** Statistical information for Fig. 2e and h. *N* values used to calculate the statistics, degrees of freedom (*df*), *F* values, *t*-values, and *p* values. The *N* values used to calculate the statistics are derived from two independent experiments.

| Fig. 3a Nuclear elasticity | | | |
| --- | --- | --- | --- |
| Group | *N* |  | |
| PC9serum- | 29 |  |  |
| PC9serum-TGFβ+ | 30 |  |  |
| Welch’s t-test | *df* | *t* | *p* |
| PC9serum- vs. PC9serum- TGFβ+ | 50.27 | 3.415 | 0.0013 |
| Fig. 3c Cell membrane elasticity | | | |
| Group | *N* |  | |
| PC9serum- | 31 |  |  |
| PC9serum-TGFβ+ | 30 |  |  |
| Welch’s one-way ANOVA | *df* | *F* | *p* |
| PC9serum- vs. PC9serum- TGFβ+ | 50.34 | 2.246 | 0.8492 |

**Supplemental Table 4 |** Statistical information for Fig. 3. *N* values used to calculate the statistics, degrees of freedom (*df*), *F* values, *t*-values, and *p* values. The *N* values used to calculate the statistics are derived from two independent experiments.

| Fig. 4b Nuclear volume | | | |
| --- | --- | --- | --- |
| Group | *N* |  | |
| PC9serum+ | 94 |  |  |
| PC9serum- | 98 |  |  |
| PC9serum-TGFβ+ | 80 |  |  |
| Welch’s one-way ANOVA | *df* | *F* | *p* |
| Factor (Group) | 2 | 129.21 | 0.000000000000000 |
| Residual | 269 |  |  |
| Games-Howell test | *df* | *t* | *p* |
| PC9serum+ vs. PC9serum- | 141.33 | 13.50 | 0.000000000000000 |
| PC9serum- vs. PC9serum-TGFβ+ | 156.91 | 0.91 | 0.64 |
| PC9serum+ vs. PC9serum-TGFβ+ | 158.48 | 12.00 | 0.000000000000019 |
| Fig. 4c Nuclear circularity | | | |
| Group | *N* |  | |
| PC9serum+ | 99 |  |  |
| PC9serum- | 98 |  |  |
| PC9serum-TGFβ+ | 80 |  |  |
| Welch’s one-way ANOVA | *df* | *F* | *p* |
| Factor (Group) | 2 | 2.86 | 0.059 |
| Residual | 274 |  |  |
| Games-Howell test | *df* | *t* | *p* |
| PC9serum+ vs. PC9serum- | 191.96 | 1.45 | 0.32 |
| PC9serum- vs. PC9serum-TGFβ+ | 176.00 | 0.93 | 0.62 |
| PC9serum+ vs. PC9serum-TGFβ+ | 174.27 | 2.52 | 0.034 |
| Fig. 4e Cell adhesion area | | | |
| Group | *N* |  | |
| PC9serum+ | 36 |  |  |
| PC9serum- | 54 |  |  |
| PC9serum-TGFβ+ | 51 |  |  |
| Welch’s one-way ANOVA | *df* | *F* | *p* |
| Factor (Group) | 2 | 6.76 | 0.0016 |
| Residual | 138 |  |  |
| Games-Howell test | *df* | *t* | *p* |
| PC9serum+ vs. PC9serum- | 44.22 | 2.64 | 0.030 |
| PC9serum- vs. PC9serum-TGFβ+ | 102.95 | 3.78 | 0.00076 |
| PC9serum+ vs. PC9serum-TGFβ+ | 43.31 | 0.88 | 0.66 |

**Supplemental Table 5 |** Statistical information for Fig. 4. *N* values used to calculate the statistics, degrees of freedom (*df*), *F* values, *t*-values, and *p* values. The *N* values used to calculate the statistics are derived from two independent experiments.

**SI References**

1 Hayashi-Takanaka Y. *et al.* Distribution of histone H4 modifications as revealed by a panelof specific monoclonal antibodies. *Chromosome Res* (2015) **23**, 753–766 (2015). <https://doi.org/10.1007/s10577-015-9486-4>
